# Phylogenetic analysis and in silico studies link spike Q675H mutation to SARS-CoV-2 adaptive evolution

**DOI:** 10.1101/2021.10.27.466055

**Authors:** Bertelli Anna, D’Ursi Pasqualina, Campisi Giovanni, Messali Serena, Milanesi Maria, Giovanetti Marta, Ciccozzi Massimo, Caccuri Francesca, Caruso Arnaldo

**Affiliations:** Section of Microbiology, Department of Molecular and Translational Medicine, University of Brescia, 25123 Brescia, Italy; Institute of Technologies in Biomedicine, National Research Council, 20090 Segrate, Italy; Section of Experimental Oncology and Immunology, Department of Molecular and Translational Medicine, University of Brescia, 25123 Brescia, Italy; Laboratório de Flavivírus, Instituto Oswaldo Cruz, Fundação Oswaldo Cruz, Rio de Janeiro, Brazil; Medical Statistics and Molecular Epidemiology Unit, University Campus Bio-Medico of Rome, Rome, Italy

**Keywords:** SARS-CoV-2, VOC, Q675H spike mutation, Furin cleavage, Phylogenesis, Molecular dynamics

## Abstract

Genotype screening was implemented in Italy and showed a significant prevalence of new SARS-CoV-2 mutants carrying Q675H mutation, near the furin cleavage site of spike protein. Currently, this mutation, which is expressed on different SARS-CoV-2 lineages circulating worldwide, has not been thoughtfully investigated. Therefore, we performed phylogenetic and biocomputational analysis to better understand SARS-CoV-2 Q675H mutants’ evolutionary relationships with other circulating lineages and Q675H function in its molecular context. Our studies reveal that Q675H spike mutation is the result of parallel evolution because it arose independently in separate evolutionary clades. *In silico* data show that the Q675H mutation gives rise to a hydrogen-bonds network in the spike polar region delimiting the conformational space of the highly flexible loop containing the furin cleavage site. This results in an optimized directionality of arginine residues involved in interaction of spike with the furin binding pocket, thus improving proteolytic exposure of the viral protein. Furin was found to have a greater affinity for Q675H than Q675 substrate conformations. As a consequence, Q675H mutation is likely to confer a fitness advantage to SARS-CoV-2 by promoting a more efficient viral entry. Interestingly, here we show an ongoing increase in the occurrence of Q675H spike mutation in the most common SARS-CoV-2 variants of concern (VOC). This finding highlights that, VOC are still evolving and start acquiring the Q675H mutation. At the same time, it suggests that our hypothesis of fitness advantage prompted by Q675H could be concrete.

## 1. Introduction

Severe acute respiratory syndrome-related coronavirus 2 (SARS-CoV-2) emerged in December 2019 in Wuhan, China, and then spread all around the world causing coro-navirus disease 19 (COVID-19) pandemic [1]. In October 2020, after the first pandemic wave, United Kingdom faced a rapid rise in COVID-19 positive cases. This prompted deep SARS-CoV-2 genomic sequencing which revealed that most of the viral sequences belonged to a new single phylogenetic cluster and led to the identification of a new SARS-CoV-2 lineage, defined as B.1.1.7. By the end of 2020, this new lineage became predominant in United Kingdom and it was defined as a variant of concern (VOC) 202012/01 [2, 3]. Approximately in the same period, in South Africa, Brazil and United States of America new circulating lineages were described for the first time, respec-tively designated as B.1.351 [4], P1 descending from B.1.1.28 lineage [5] and B.1.427/B.1.429 [6]. These last variants were also named as VOC because their poly-morphisms enhance viral transmissibility and may enable the viral escape from vac-cine-elicited neutralizing antibodies. All the VOC are characterized by a typical mutational pattern, in which most of these mutations are located in the spike receptor binding domain (RBD), the most variable part of the coronavirus genome [7, 8], lead-ing to an increased affinity to human angiotensin-converting enzyme 2 (ACE2) and consequently, to enhanced infectivity [9, 10]. Indeed, these new lineages are becoming predominant not only in the countries where they emerged but also in the neighboring ones. Thus, these findings suggest that viral evolution is ongoing and aims to adapt to the host and to acquire more fitness.

SARS-CoV-2 entry requires sequential cleavage of the spike glycoprotein at the S1/S2 and the S2’ cleavage sites to mediate membrane fusion [11–13]. A notable feature of SARS-CoV-2 is a polybasic site (PRRAR) at the junction of S1 and S2 [14], that consti-tutes a furin binding site for proteolytic cleavage [15]. This polybasic site is absent from the SARS-CoV-2 closest lineage B betacoronaviruses [16, 17], although similar pol-ybasic cleavage sites are found in more distantly related coronaviruses such as HKU1 (RRKRR) and OC43 (RRSR) [18, 19]. The functional consequence of the polybasic cleavage site in SARS-CoV-2 spike is not completely understood, but it is likely to pro-vide the virus with its unique cross-species transmissibility and serve as a pathogenet-ic element in human infection [13, 20]. In avian influenza virus, acquisition of polyba-sic cleavage sites in hemagglutinin is known to convert low-into high-pathogenicity viruses [21] and to facilitate infection of a wide variety of cell types [21–23]. At the same time, there is evidence that promoting a more efficient cleavage of spike enables MERS-like coronaviruses from bats to infect human cells [24]. Interestingly, insertion of a furin cleavage site at the S1-S2 junction of SARS-CoV spike enhances viral infec-tivity [25]. More recently, Peacock et al. [26] showed that the polybasic cleavage site endows SARS-CoV-2 with a selective advantage in lung cells and in primary airway epithelial cells, by enhancing cell entry of progeny virions.

Despite acquisition of the polybasic cleavage site is a notable feature of SARS-CoV-2, it remains suboptimal for the enzymatic cleavage exerted by furin [15]. Therefore, it is possible that further optimization of this site could result in higher human-to-human transmissibility and possibly in widening SARS-CoV-2 host range [27]. If this hypothe-sis is correct, we have to expect the emergence of rapidly spreading variants displaying mutations within and/or proximal to the S1/S2 cleavage site predicted to enhance furin binding and proteolytic activity.

Furin cleavage site motif includes about 20 amino acid residues defining a core region (8 amino acids, positions P6-P2’) packed inside the furin binding pocket, and two polar regions (8 amino acids, positions P7-P14’; and 4 amino acids, positions P3’-P6’) located outside the furin binding pocket. The physical properties of the core region contribute to the binding strength of the furin substrate to the enzyme, while the polar regions fa-cilitate the accessibility of the core region to the furin binding pocket [28]. Here we de-scribe the emergence of SARS-CoV-2 strains carrying mutation involving the substitu-tion of a glutamine (Q) with a histidine (H) residue at position 675 (Q675H), within the polar region of spike proximal to the S1/S2 furin cleavage site. We also demonstrate that Q675H mutation contributes to form a hydrogen bonds (H-bonds) network that facilitates the accessibility of the spike core region to the furin binding pocket and im-proves the orientation of arginine residues to increase furin binding strength to the vi-ral substrate. Moreover, we highlight that Q675H mutation is the result of a process of parallel evolution, and that multiple SARS-CoV-2 VOC displaying the Q675H muta-tion have recently emerged. Altogether, our results point to a role of Q675H spike mu-tation in enhancing viral fitness by ensuring a more efficient spike proteolytic cleav-age.

## 2. Materials and Methods

### Detection of SARS-CoV-2

Nasopharyngeal specimens were collected from the end of January to the end of March 2021 at the Brescia Civic Hospital, (Brescia, Lombardy, Italy), using FLOQSwabs in the universal transport medium (UTM) (COPAN, Brescia, Italy). Viral RNA was extracted from 300 μl of UTM with Nimbus automatic system (Arrow Diagnostics, Genoa, Italy), according to the manufacturer’s instructions. Amplification was performed on BioRad CFX PCR machine (Bio-Rad Laboratories S.r.l., Milan, Italy) using the Allplex™ 2019-nCoV Assay (Seegene Inc. Seoul, Korea). Ct values were automatically calculated using the 2019-CoV Viewer analysis software (Seegene).

### Viral RNA extraction and amplification

Total RNA was extracted from 200 μl of PCR-positive nasopharyngeal swabs using QIAamp DSP Virus Kit® (Qiagen, Hilden, Germany) according to the manufacturer’s instructions. RNA was eluted in 30 μl AVE, following manufacturer’s guidelines and stored at −80 °C until use. SARS-CoV-2 RNA was reverse-transcribed and PCR ampli-fied using SuperScript™ III One-Step RT-PCR System with Platinum™ Taq DNA Polymerase (Thermo Fisher Scientific, Carlsbad, CA, USA) in a 50 μl reaction contain-ing 25 μl of reaction mix, 9 μl of MgSO4, 2 μl of SuperScriptTM III RT/PlatinumTM Taq Mix, 0.2 μM of sense and antisense primers, and 12 μl of extracted RNA. The amplifi-cation conditions were as follows: 50° C for 30 min (for reverse transcription) and 94 °C for 2 min for Taq DNA polymerase activation, followed by 40 cycles (94°C 15 sec, 56°C 30 sec, 68°C 2 min) and a final cycle at 68°C 7 min. PCR primers used in the reaction were: SARS2-S-F5, 5’-GATGAAGTCAGACAAATCGCTCCAGG and SARS2-S-R6, 5’-TTCTGCACCAAGTGACATAGTGTAGGCA. Afterwards, PCR product were checked on a 1% agarose gel and were purifying through QIAquick PCR Purification Kit® (Qiagen, Hilden, Germany) according to the manufacturer’s instructions and quantified using the Qubit DNA HS Assay Kit (Thermo Fisher Scientific, Carlsbad, CA, USA). To assess the occurrence of mutations of interest in Sars-CoV-2 sample, purified PCR products were sequenced using the SARS2-S-F5 and SARS2-S-R6 primers with the BigDye terminator v3.1 cycle sequencing kit on SeqStudio Genetic Analyzer (Thermo Fisher Scientific, Carlsbad, CA, USA). The derived sequences were analyzed with Ge-neious software (version 11.1.5) (Biomatters Ltd, New Zealand), using the sequence NC_045512.2 as SARS-CoV-2 reference.

### Whole genome sequencing

Virus genomes were generated using Paragon Genomics’ CleanPlex multiplex PCR Research and Surveillance Panel, according to the manufacturer’s protocol [29–31]. This panel works with an amplicon-based approach, targeting 343 partially overlap-ping subgenomic region that cover the entire SARS-CoV-2 genome. Briefly, a 10 μl multiplex PCR was performed with two pooled primer mixtures; the cDNA was reverse transcribed with random primers and it was used as a template. After ten rounds of amplification, the reaction was terminated by the addition of 2 μl of stop buffer and then the two PCR products were pooled and purified. The resulting solution was treated with 2 μl of CleanPlex digestion reagent at 37°C for 10 min to remove non-specific PCR products. After a magnetic bead purification, the PCR product was subjected to further 25 rounds of amplification in a secondary PCR with a pair of pri-mers to produce the metagenomic library. Subsequently, purified libraries were quan-tified using the Qubit DNA HS Assay Kit (Thermo Fisher Scientific). Then, libraries were adjusted to 4 nM concentration and finally equal amounts of each library were pooled. Library pool was denatured with NaOH and further diluted to 8 pM. Pooled libraries were subsequently loaded in a 300-cycle sequencing cartridge and deep se-quencing was performed on an Illumina MiSeq platform. Sequencing raw data were checked for quality using FastQC (https://www.bioinformatics.babraham.ac.uk/projects/fastqc/) and then analyzed with the specifically designed software SOPHiA GENETICS’ SARS-CoV-2 Panel (SOPHiA GENETICS, Lausanne, Switzerland).

### Phylogenetic analysis

The 11 whole genome sequences of SARS-CoV-2, from Brescia, northern Italy, reported in this study, were first evaluated using the Phylogenetic Assignment of Named Global Outbreak LINeages tool, available at https://github.com/hCoV2019/pangolin [32], for the lineages assesment. Newly whole genome sequences from this study were then aligned with European SARS-CoV-2 genomes strains available in GISAID from March 2020 to March 2021. Low-quality genomes and nearly identical sequences (genetic similarity>99.99%) were excluded, obtaining a final dataset of 1487 European se-quences. The same principle was then applied to obtain the four additional representa-tive datasets of the circulating VOCs (alpha, beta, gamma and delta) generated by us-ing downsampled worldwide SARS-CoV-2 genomes available in GISAID from No-vember 2020 to July 2021. In particular, the B.1.1.7 dataset includes n=1500 genomes plus n=278 strains carrying the Q675H mutation; the B.1.351 dataset includes n=1100 genomes plus n=104 strains carrying the Q675H mutation; the P.1 dataset includes n=1400 genomes plus n=59 carrying the Q675H mutation; the B.1.617+ A.Y.x dataset includes n=1776 genomes plus n=664 carrying the Q675H mutation. Whole genome sequences were aligned with MAFFT (FF-NS-2 algorithm) using default parameters [33]. The alignment was manually curated with Aliview [34] to remove artifacts at the ends and within the alignment. Phylogenetic analysis was performed using IQ-TREE-2 under the best fit model according to Bayesian Information Criterion (BIC) indicated by the Model Finder application implemented in IQ-TREE-2 [35].

### Biocomputational studies of Q675H variants

Structure of SARS-CoV-2 spike protein, model 6vxx_1_2_1 (aa 1-1146) without gly-cans, were taken from the CHARMM-GUI website. (http://www.charmm-gui.org/docs/archive/covid-19). Pymol mutagenesis wizard was used to model Q675H and D614G for both wild-type (wt) and mutant systems [36]. The DynaMut [37], webserver was used to predict the effect of genetic variants on the sta-bility and flexibility of Q675H mutant spike protein. Molecular dynamics simulations were performed for the SARS-CoV-2 spike proteins to estimate the stability and dy-namics features of Q675 and Q675H. For this purpose, we used Amber 18 [38] package and the ff14SB force field parameters for protein. Each complex was solvated in a pe-riodic cubic water box using the TIP3P water model with 10 Å between the solutes and the edges of the box, then a suitable number of Na+ and Cl- ions was added to neutral-ize the whole system and mimicked a salt solution concentration of 0.15M. Each sys-tem was energy minimized in 14 consecutive minimization steps, each of 2500 steps of steepest descent followed by 2500 steps of conjugate gradient, with decreasing positional restraints from 1000 to 0 kcal/mol A2 on all the atoms of the systems excluding hydrogens, with a cutoff for non-bonded interactions of 8 Å. The systems were then subjected to two consecutive steps of heating, each of 100000 steps, from 0.1 to 100 K and from 100 to 310 K in an NVT ensemble with a Langevin thermostat. Bonds in-volving hydrogen atoms were constrained with the SHAKE algorithm and 2 fs time step was used. The systems were then equilibrated at 310 K in 4 consecutive steps of 2.5 ns each in the NPT ensemble with random velocities assigned at the beginning of each step. MDs were performed over 100 ns using the pmemd CUDA program of the Amber18 package and a server Tesla K20 Graphical Processing Unit. The trajectories were analyzed to compare and observe the structural deviation between wt and mutant structures. Conformational ensembles of spike chains along the trajectories of MDs were analyzed in the differential geometry study using FleXgeo [39]. Further-more, the conformational ensembles along the trajectory were clustered by CPPTRAJ using hierarchical agglomerative algorithm with epsilon 3.0. Hydrogen bonds analysis were carried out using CPPTRAJ obtaining the hydrogen bond frequency for each res-idue pair during the molecular dynamics. The most representative conformation of each chain was used for the docking-docking protein by Cluspro webserver [40] and binding affinities studies by PRODIGY webserver [41].

## 3. Results

### Reporting of SARS-CoV-2 Q675H mutation

A rapid rise in SARS-CoV-2 B.1.1.7 lineage in Europe prompted us to implement a ge-nomic surveillance program in the Brescia province. From the end of January 2021 to the end of March 2021, we sequenced by Sanger 228 samples from SARS-CoV-2 posi-tive patients with Cycle threshold (Ct) values < 30 (Ct range = 14-27,5; Ct median = 17,5). This analysis revealed the presence of B.1.1.7 lineage from United Kingdom, B.1.351 lineage from South Africa, B.1.525 from Nigeria, and wild-type sequences. In the wild-type sequences, the number of those displaying a Q675H mutation in the spike pocket for furin binding, was considerably noteworthy. Indeed, in the considered interval of time, Q675H mutants represented 40% of all the analyzed wild-type se-quences, leading us to hypothesize a role of this mutation in SARS-CoV-2 adaptive evolution.

In order to understand if the Q675H mutation had already occurred in Italy before our observation in Brescia, a retrospective analysis of genomes displaying this mutation was retrieved in GISAID. As shown in Figure 1, in Italy, SARS-CoV-2 sequences carry-ing the Q675H mutation were uncovered for the first time in October 2020. Since then, Q675H prevalence arose until December 2020, when it reached the highest occurrence and, afterwards, a steady state of Q675H mutation rate was observed until the end of February 2021. The prevalence of this mutation started to decrease in March 2021 be-cause of the progressive establishment of the newly circulating B.1.1.7 lineage, that became predominant in Italy on any other previously circulating lineage. To note, when the B.1.1.7 lineage firstly emerged did not carry the Q675H spike mutation.

**Figure 1.**
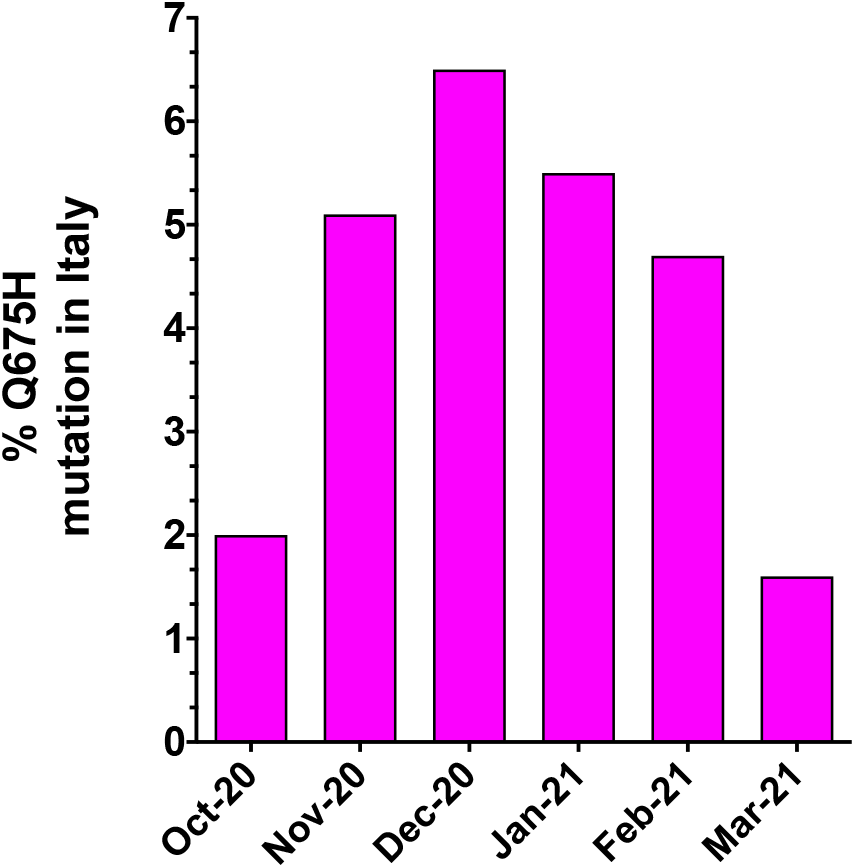
Prevalence of SARS-CoV-2 Q675H mutants in Italy from October 2020 to March 2021. The number of SARS-CoV-2 Q675H mutants has been normalized to the total number of the Italian SARS-CoV-2 sequences retrieved in GISAID.

### Phylogenetic analysis of SARS-CoV-2 sequences carrying Q675H mutation

Whole genome sequencing was performed successfully in our laboratory for 11 SARS-CoV-2 Q675H mutants. We found that Q675H mutation was due to a transition or transposition at nucleotide position 23588 (respectively, G->C or G->T). Remarka-bly, 5 out of 11 sequences belonging to B.1 lineage carried the transition G->C, whereas 6 out of 11 belonging to B.1.177 lineage carried the transposition G->T. In order to ac-curately determine evolutionary relationships of the 11 SARS-CoV-2 Q675H mutants detected in Brescia within a similar European context, a maximum likelihood (ML) tree was employed. The dataset utilized in this analysis consists of European sequences collected from the beginning of pandemic to March 2021, and belonging to different lineages. As shown in Figure 2, 5 sequences out of 11 displaying Q675H mutation we obtained gave rise to an independent cluster in the B.1 lineage. The 6 remaining clus-tered in the B.1.177 lineage with other European Q675H sequences, but scattered across the phylogenetic tree. It is worth noting that Q675H sequences represented in the phylogenetic tree cluster in different clades, in accordance with their lineage of be-longing. This finding attests for the emergence of Q675H mutation by homoplasy, a process of parallel evolution [42], in which different populations, in different coun-tries, have acquired the same advantageous genome mutations, multiple times, inde-pendently and in separate evolutionary clades [43, 44].

**Figure 2.**
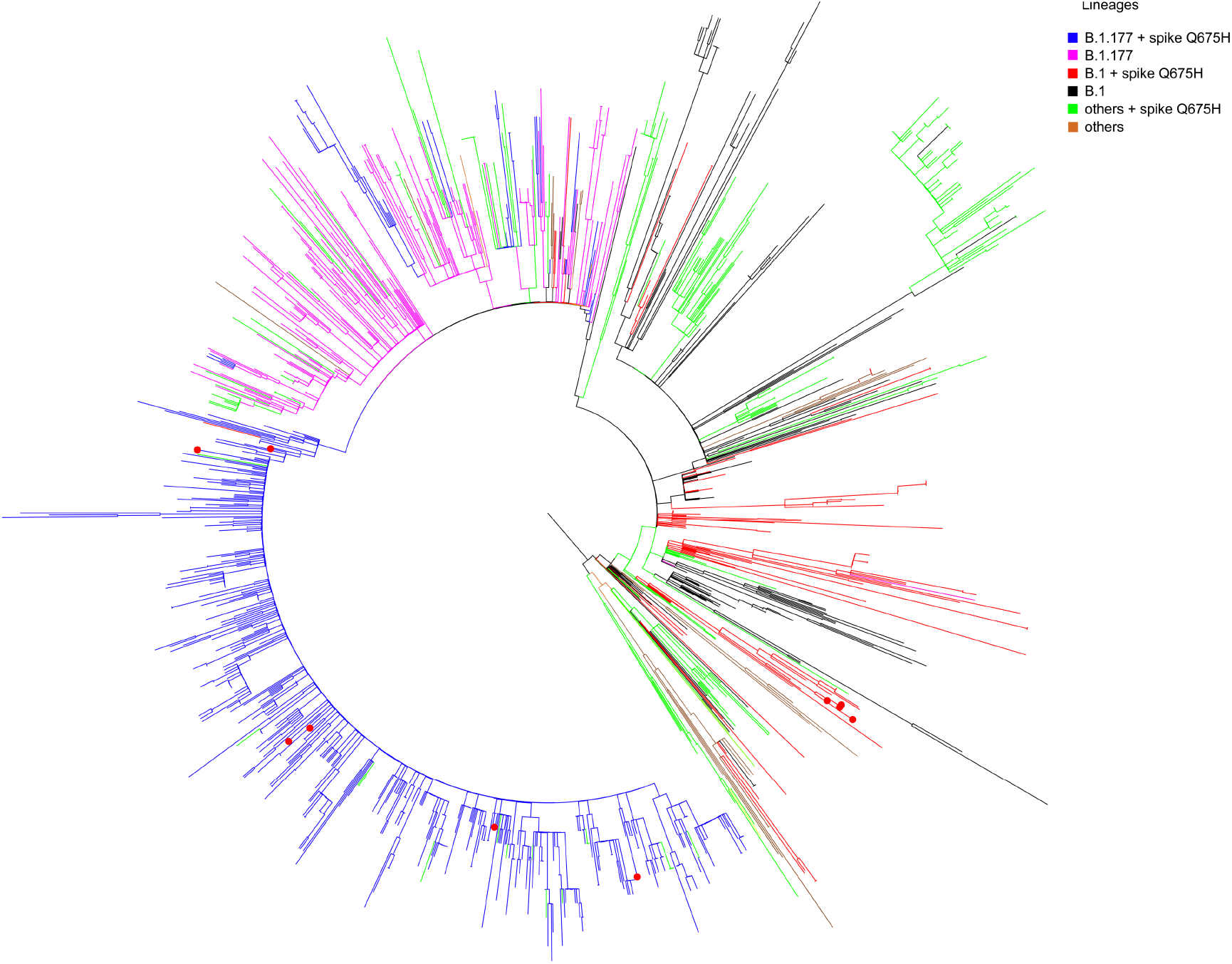
Maximum-likelihood tree of European SARS-CoV-2 sequences. The mid-point rooted, ML tree includes 1487 SARS-CoV-2 sequences retrieved in Europe from GISAID database from March 2020 to March 2021 and 11 SARS-CoV-2 spike Q675H sequences obtained in Brescia between the end of January 2021 and the end of February 2021. Branches are colored by lineages: B.1 + spike Q675H in red; B.1 in black; B.1.177 + spike Q675H in blue; B.1.177 in fuchsia; all of the other lineages carrying the Q675H mutation indicated as “others + spike Q675H” in green; all the other lineages not carrying the Q675H mutation indicated as “others” in brown. Red dots represent the 11 Brescia SARS-CoV-2 spike Q675H mutants.

### Occurrence of Q675H mutation in different VOC

In Brescia province we experienced in March 2021 the rapid spreading of SARS-CoV-2 B.1.1.7 lineage, that overtook all the others with a prevalence in April 2021 of 95,1% over the B.1.525 (3,7%) and the B.1/B.1.177 wild-type lineages (1,2%). Therefore, it is likely to assume that the gain of fitness of wild-type lineages prompted by Q675H spike mutation itself had to be modest compared to mutations in RBD displayed by B.1.1.7 lineage, that are known to confer a higher binding affinity of spike to ACE2 and in-crease virus infectivity [9, 10]. To attest its role in virus adaptive evolution, we ana-lyzed worldwide the occurrence of Q675H mutation in SARS-CoV-2 VOC through GISAID available data (https://www.gisaid.org/). When VOC firstly emerged, neither of them carried this mutation. However, as shown in Figure 3, it is remarkable to note that all the VOC are acquiring the Q675H mutation and the percentage of VOC carry-ing Q675H mutation is continuously increasing over time, even in the most recently emerged B.1.617.2 lineage, with the first appearance on June 2021. Moreover, as ex-pected, sequences displaying Q675H mutation were scattered across the phylogenetic tree of each VOC (Figure 4). This finding further confirms that Q675H mutation arises in phylogenetically distant VOC by a homoplasy event, and underlines that the hy-pothesis of a fitness advantage prompted by Q675H mutation can be concrete.

**Figure 3.**
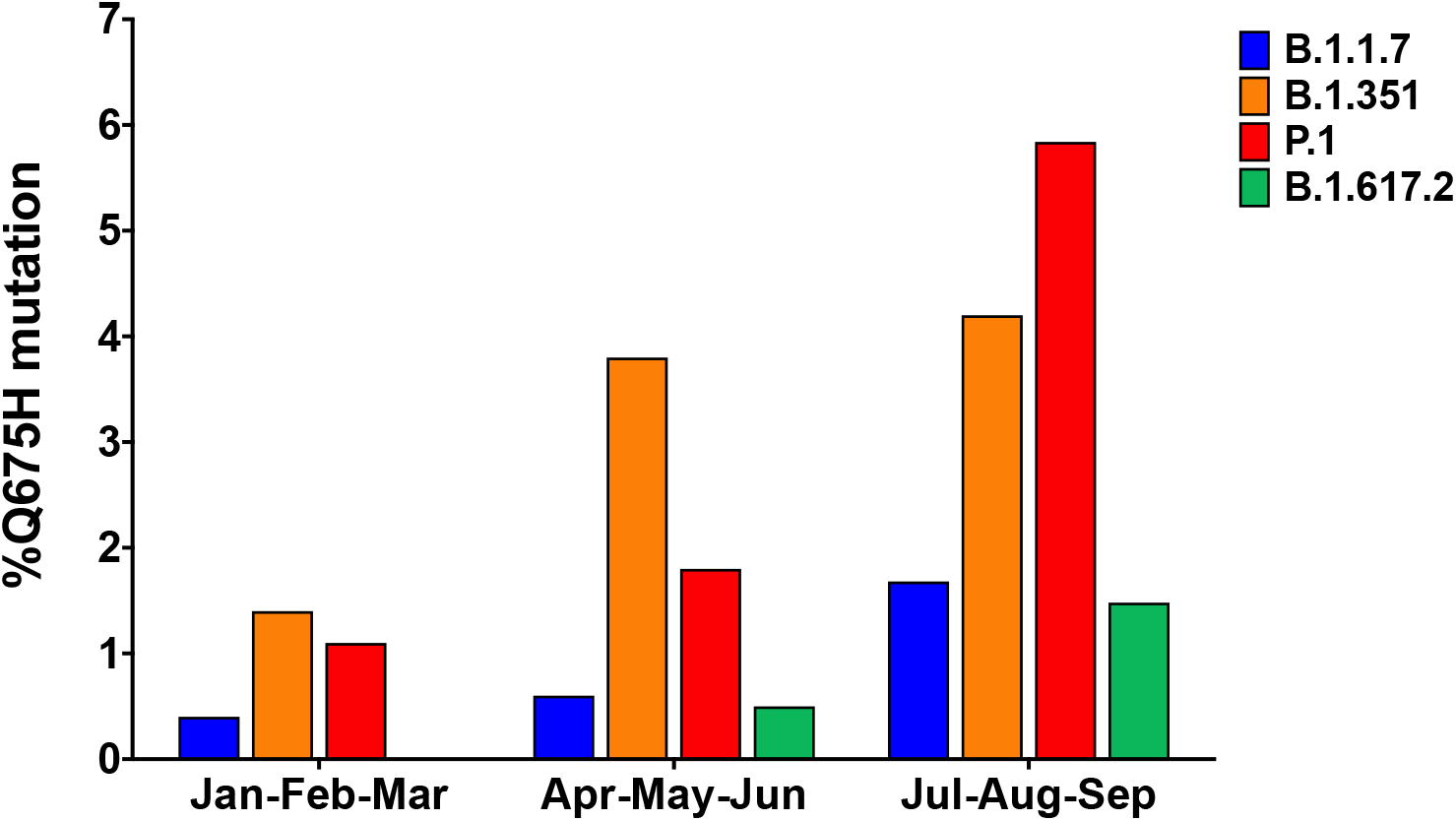
Prevalence of Q675H spike mutation in VOC from January 2021 to September 2021. The number of SARS-CoV-2 Q675H mutants VOC has been normalized to the total number of SARS-CoV-2 VOC sequenced globally.

**Figure 4.**
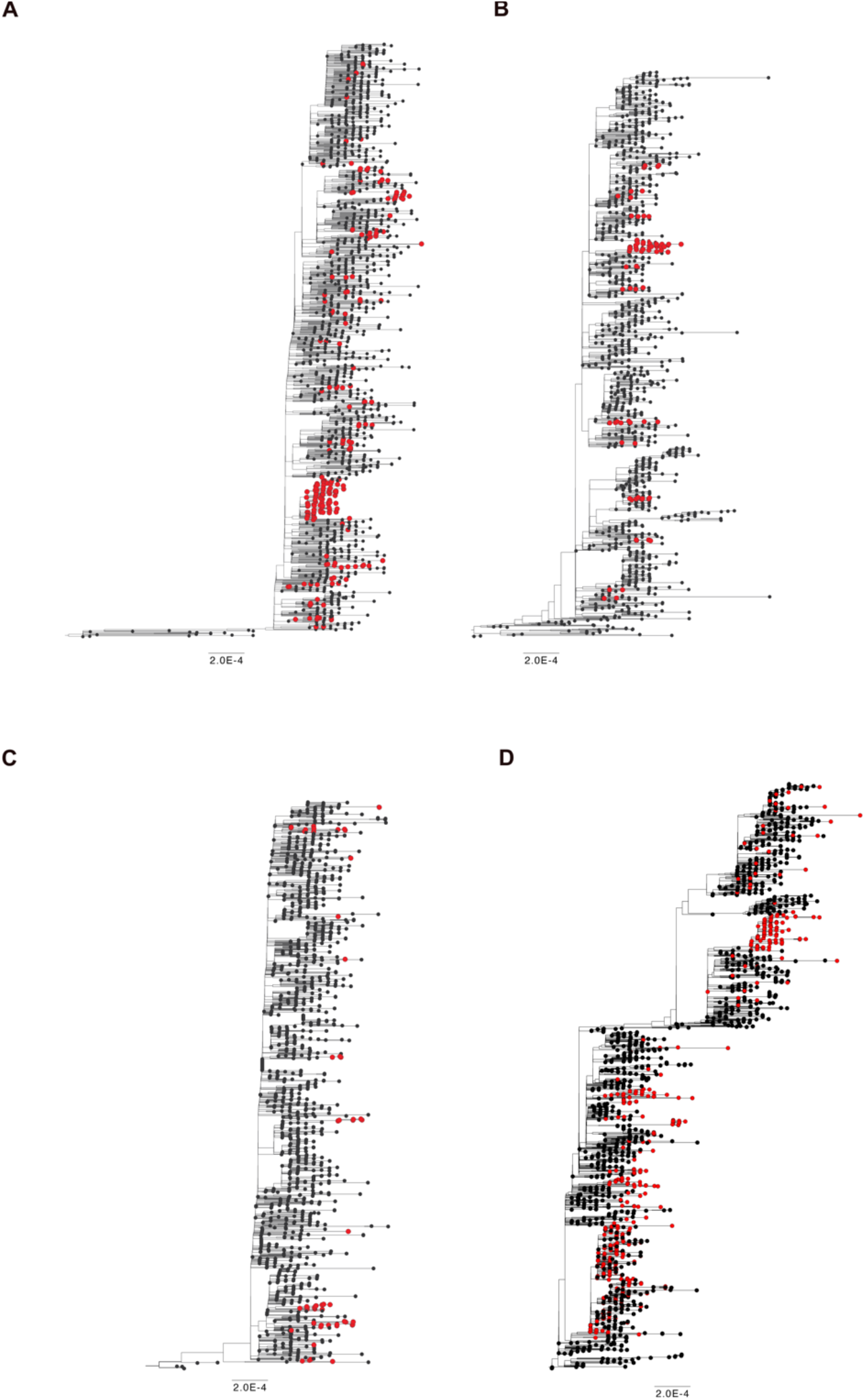
Time-scaled maximum likelihood trees including sequences representative of worldwide SARS-CoV-2 VOC (black circles) from November 2020 until July 2021. Genomes carrying the mutation Q675H are highlighted with red circles. (A) B.1.1.7 lineage includes 1500 genomes plus 278 carrying Q675H mutation; (B) B.1.351 lineage includes 1100 genomes plus 104 carrying Q675H mutation; (C) P.1 lineage includes 1400 genomes plus 59 carrying Q675H mutation; B.1.617 + A.Y.x lineage includes 1776 genomes plus 664 carrying the Q675H mutation.

### Q675H confers a lower structural variability to the furin cleavage site loop by form-ing a distinct H-bond network

Based on our findings, we hypothesize that SARS-CoV-2 Q675H spike mutation may enhance viral fitness and host adaptation. Biocomputational studies were performed to explore the role of Q675H mutation in the context of the furin binding pocket. The Q675H mutation is located in the spike protein adjacent to the polybasic S1/S2 furin cleavage site 682-RRAR*S −686. The stability of the spike tertiary structure was evaluated using DynaMut web server [37]. Q675H mutation showed a positive (ΔΔG value is 0.665Kcal Kcal/mol) and a negative (ΔΔSVibENCoM value is –0.108 kcal.mol-1K-1) change in vibrational entropy energy between wild-type and mutant proteins. A ΔΔG above zero and a ΔΔSVibENCoM below zero represent protein stabilization and stiff-ening of the protein structure, respectively.

To further characterize the structural stability and dynamic features of spike express-ing Q675 or Q675H, we performed 100ns molecular dynamics (MDs) simulations starting from the 6VXX charm-gui model (aa 1-1146) without glycans. Conformational ensembles of spike chains along the trajectories of MDs were analyzed representing protein backbone as a 3D regular curve, and characterized by curvature and torsion values per residue. We used the differential geometry (DG)-based representation of the protein structure [39] because it is optimal for backbone protein flexibility analysis and particularly advantageous for highly flexible regions of protein, such as the SARS-CoV-2 spike protein loop containing the furin cleavage site. The visual inspec-tion of the curvature (κ) and torsion (τ) values observed on the furin cleavage site loop shows that compared to Q675 chains (Figure 5A-B-C), a lower structural variability in spanning residues ranging from 675 to 683 in the Q675H chains was observed (Figure 5D-E-F). Further analysis showed that Q675H forms a distinct H-bond network in-volving residues at position 676, 677, 678 and 679 that are localized near the Q675H residue (Figure 5).

**Figure 5.**
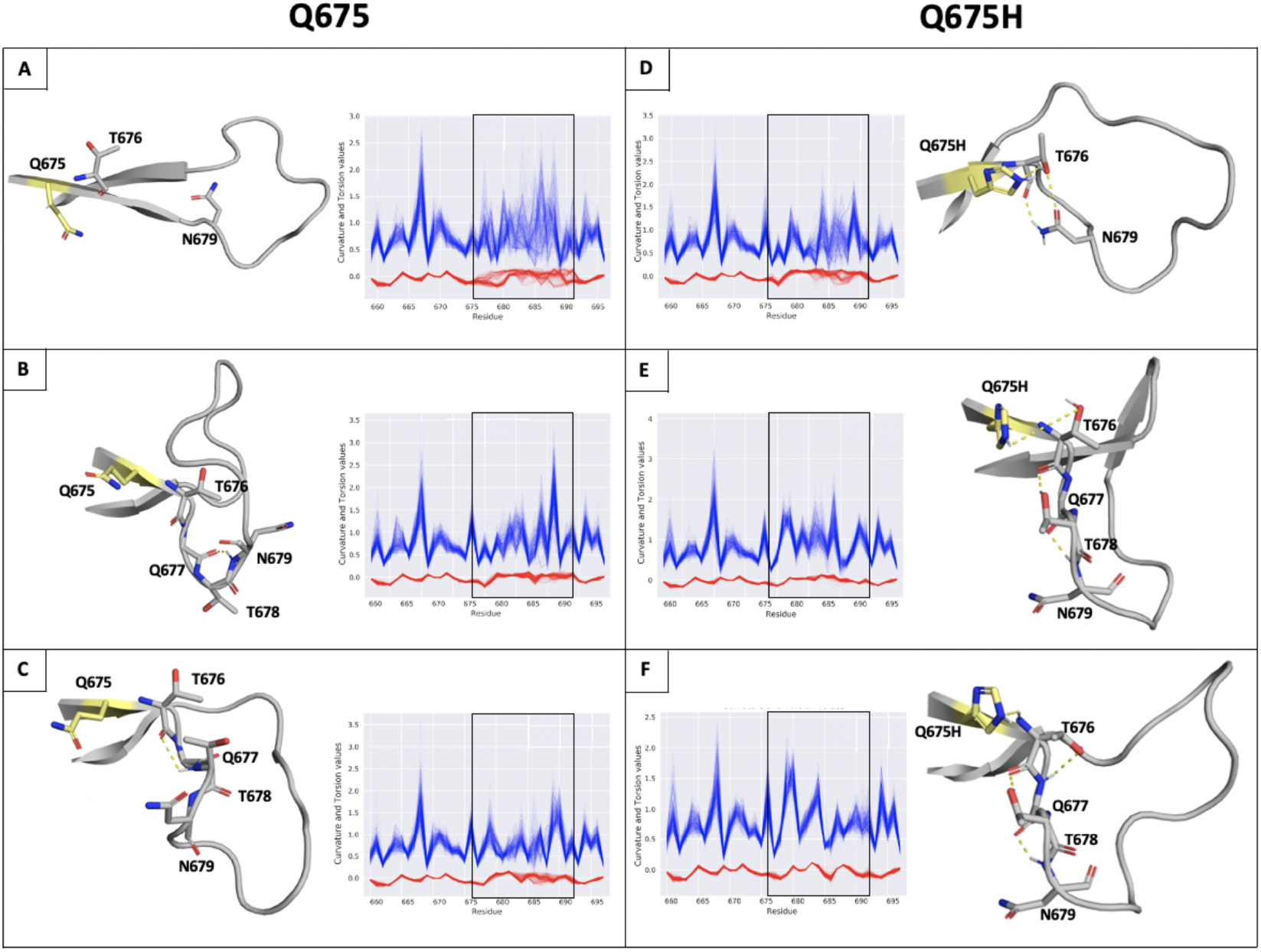
(Panel A-F) H-bonds and conformational flexibility around furin cleavage site loop. Representative conformation of furin cleavage site loop for each chain of Q675 (left) and Q675H (right) obtained by cluster analysis. The residues involved in H-bonds network are shown in ribbon and sticks respectively. Chains A, B and C of Q675 are shown in panels A, B and C; chains A, B, and C of Q675H are shown in panels D, E and F. The H-bonds are shown as dotted lines. Plot of curvature (blue line) and torsion (red line) of the Q675 and Q675H of ensemble for each chain in function of sequence for residues 656 to 697 around furin cleavage site loop are shown. The curvature and torsion values of the residues of furin cleavage site affected by hydrogen bond network is highlighted (window).

### Analysis of substrate conformations within the furin binding pocket

The H-bond network is located outside the furin binding pocket and could facilitate the accessibility of the spike core region to the furin binding pocket [28]. In order to evalu-ate the capability of our models to interact with the furin binding pocket in the correct orientation, the conformational ensembles along the trajectory were clustered. Then the side chain of R685 in position P1, R683 in position P3 and R682 in position P4 of the furin recognition motif were evaluated on the basis of their spatial arrangement at and around P1 and P4 residues of the furin inhibitor co-crystallized in the furin binding pocket. The three arginine residues were analyzed in every chain of the trimeric spike protein, according to their optimal orientation into the enzyme pocket (Figure 6A). For each chain the representative conformation of the most-populated cluster was selected to study the interaction between spike and furin proteins by the ClusPro web server for protein-protein docking [40]. The poses complexes obtained from the docking simula-tions were analyzed by visual inspection and filtered evaluating orientation of the ar-ginine side chains compared with the orientation of P1 and P4 residues of the furin in-hibitor and the lowest ClusPro score. As shown in Figure 6B, three active substrate conformations for Q675H and two for Q675 were selected considering arginines within the 682-RRAR-685 cleavage site: P1-S1 substrate conformation extends only the guanidino side chain of the R685 in position P1 into the S1 pocket; P1-S1/P3-S4 sub-strate conformation puts also the guanidine side chains of the R683 in position P3 in the S4 pocket; and P1-S1/P4-S4 substrate conformation extends R685 in position P1 and R682 in position P4 into the S1 and S4 pocket, respectively. The P1-S1/P4-S4 sub-strate conformation was observed only for Q675H. In Q675, the P1-S1/P3-S4 substrate conformation arranged the three arginine chains in a non-alternating way, therefore the pocket S4 can be occupied by R683 in position P3 only, instead of R682 in position P4. This because R683 in position P3 is closer to R685 in position P1 and generates a steric hindrance for R682 in position P4. The steric effect prevents the interaction of the R682 in position P4 with the S4 pocket. In the Q675H, steric hindrance among the arginines does not occur because of a y–conformation which arranges the three ar-ginines alternately and puts the two arginines of the P1-S1/P3-S4 and the P1-S1/P4-S4 substrate conformation at the two arms of the y conformation, thus orienting them in the same direction. This means that the two arginine residues involved in the interac-tion with the enzymatic site of furin are not affected by the steric hindrance operated by the third one. Consequently, they are able to better fit within the furin binding pocket.

**Figure 6.**
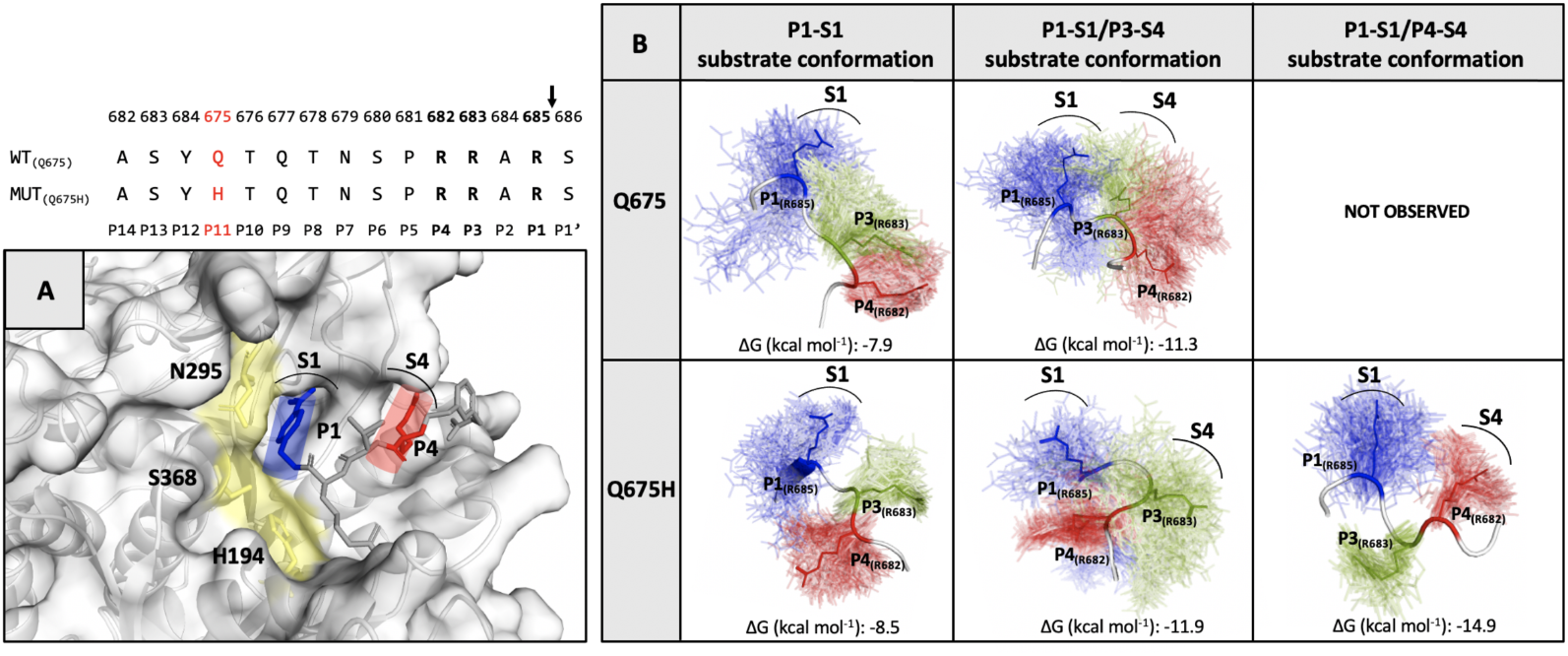
(Top) The furin cleavage site sequence in Q675 and Q675H is shown. (panel A) The X-Ray structure of the inhibitor (m-guanidinomethyl-phenylacetyl-Arg-Val-Arg-(4-amidomethyl)-benzamidine) bound to active site of furin (PDB code 5JXH). The catalytic domain of furin is shown in yellow, the locations of the P1 (blue) and P4 (red) residues of inhibitor are shown in sticks while their respective S1 and S4 interaction pockets are labelled and indicated by an arc. (panel B) The substrate conformations of the trimeric spike of Q675 and Q675H are shown. In the conformational ensemble the side chain of the three-arginine residues of the furin recognition motif in their spatial arrangement is shown in line P1-R685 (blue), P3-R683 (green) and P4-R682 (red) and the arginine residues of the most representative conformation are highlighted in sticks. The arginine residues for each substrate conformation involved in the interaction with their respective S1 and S4 furin pockets are labelled.

### Furin has greater affinity for Q675H than Q675 substrate conformations

The affinity of furin for the three substrate conformations was evaluated by calculat-ing the binding affinity of each spike furin cleavage site with the furin enzyme com-plex, using the PRODIGY web-server [41]. The binding affinity is minor for P1-S1 sub-strate conformation (−7,9 kcal mol-1 for Q675 and −8,5 Kcal mol-1 for Q675H), and in-creases for P1-S1/P3-S4 (−11,3 kcal mol-1 for Q675 and −11,9 kcal mol-1 for Q675H), and for P1-S1/P4-S4 (−14, 9 kcal mol-1 for Q675H) substrate conformations (Figure 6B). Sub-strate conformations of Q675H have a higher binding affinity than Q675. This is due to the fact that Q675H always presents a y-conformation with the three arginines alter-natively arranged. Furthermore, the substrate conformations in the most populated clusters were distributed as follows: P1-S1 substrate conformation makes up 70% (Q675) and 46% (Q675H) of the chain B; P1-S1/P3-S4 substrate conformation makes up 50% (Q675) and 30% (Q675H) of the C and A chains, respectively; and P1-S1/P4-S4 substrate conformation makes up 70% (Q675H) of the chain C. Overall Q675H has a higher number of active substrate conformations than Q675. P1-S1/P3-S4 of Q675 and P1-S1/P4-S4 of Q675H substrate conformations are those with the highest binding af-finity to furin, and Figure 7 shows their binding poses in the context of furin cleavage site.

**Figure 7.**
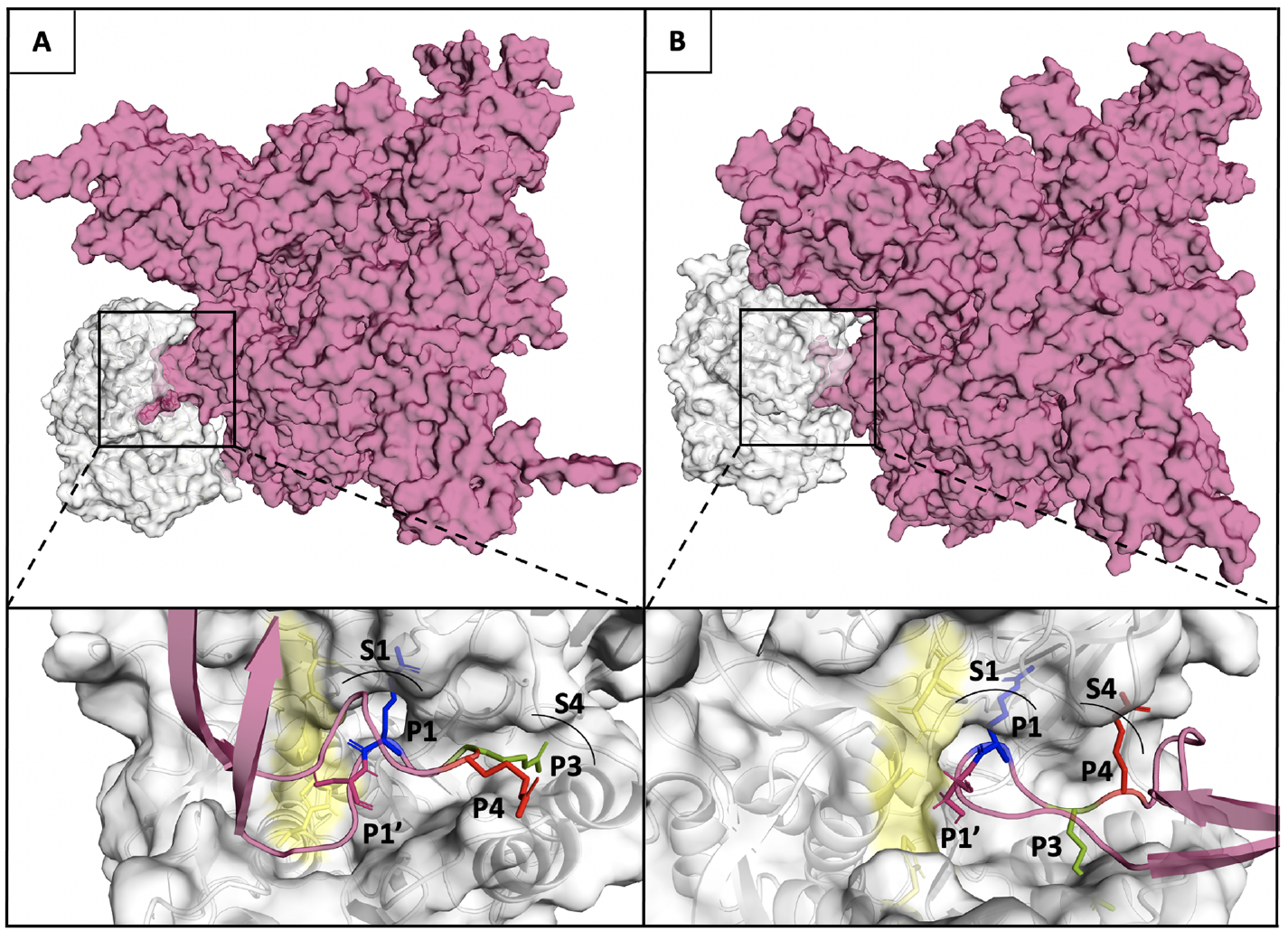
(Upper panel A and B) Furin and Q675 (left) or Q675H (right) models are shown in Connolly surface gray and fuchsia respectively. (Lower panel A and B) The binding pose of the arginines of the furin cleavage site motif in P1-S1/P3-S4 substrate conformation of Q675 (left) and P1-S1/P4-S4 substrate conformation of Q675H into the corresponding binding pockets S1 and S4 are shown. The catalytic domain of furin is shown in yellow; the locations of the P1 (blue), P3 (green) and P4 (red) of arginine involved in the interaction with enzyme are shown in sticks, while their respective S1 and S4 pockets are labelled and indicated by an arc.

### Worldwide analysis of SARS-CoV-2 lineages carrying the Q675H spike mutation

As a confirmation of our hypothesis, we analyzed Q675H spike mutation occurrence in all the SARS-CoV-2 globally circulating lineages from the beginning of the pandemic. Of note, up to October 26th 2021, 8 SARS-CoV-2 lineages (B.1.438.2, B.1.438.3, B.1.438.1, B.1.1.385, P.6, P.1.9, B.1.599, B.1.544) are documented worldwide to carry the Q675H spike mutation in the majority of their sequences retrievable in GISAID. In particular, in B.1.438.1, B.1.438.2, B.1.438.3 and B.1.1.385 lineages Q675H and D614G are the sole mutations located in the Spike protein. In B.1.438.2 and B.1.438.3 lineages Q675H is documented in the 100% of the sequences belonging to these lineages, highlighting that Q675H spike mutation is one of the characteristic mutations necessary to define these lineages. Moreover, B.1.438.1 lineage, which appeared for the first time in June 2020, is the most represented globally with 8831 sequences, 99% of which described in Canada [45].

## 4. Discussion

It has been speculated that the polybasic motif in SARS-CoV-2 spike was acquired af-ter animal-to-human transmission and it is possible that it may be not stable in the human host or undergo to further adaptation. Indeed, a panel of SARS-CoV-2 variants with various deletions directly affecting the polybasic cleavage itself, or the flanking sequence QTQTN, were identified in cultured cells and found to easily emerge in vitro [46–48]. These findings indicate that an efficient mechanism for deleting these regions exists and that they are not fixed in the virus. On the other hand, the deletion of the polybasic region and of the flanking region QTQTN is rare in vivo [47, 48], strongly suggesting that the entire region may be under selective pressure in humans [46, 49]. It is worth noting that SARS-CoV-2 variants with deletions at the S1/S2 junction of the spike protein cause no apparent disease in hamster, despite replicating in the upper respiratory tract as the wildtype virus [48]. However, unlike wild-type virus, mutants with such deletions did not cause pathological changes in the lung and did not induce elevation of proinflammatory cytokines [48]. These observations led Wang et al., [48] to suggest that the polybasic cleavage motif is a virulence element in SARS-CoV-2 and that its removal makes the virus more similar to a common cold respiratory corona-virus.

From the end of January 2021 to the end of March 2021, the genomic surveillance pro-gram implemented by our laboratory in Brescia, Italy, led us to detect an increased number of SARS-CoV-2 wild-type sequences displaying a Q675H mutation within the polar region of the spike pocket for furin binding. In particular, we found Q675H mu-tants detected in our laboratory belonged to B.1 and B.1.177 lineages, and in the considered interval of time, they represented 40% of all the wild-type SARS-CoV-2 ana-lyzed sequences. Our data was in agreement with a general increase of this mutation in different lineages circulating worldwide. Phylogenetic analyses attested that Q675H mutation occurred by a homoplasy event, leading us to assume that it was aimed to enhance viral fitness to the human host.

On March 2021 we observed in Italy a dramatic decline of the Q675H mutation rate, because of the progressive establishment of B.1.1.7 lineage over any other circulating wild-type lineage. At the time of its emergence, the B.1.1.7 lineage did not display the Q675H mutation. However, here we underline the finding that B.1.1.7 lineage as well as other VOC are starting to acquire this mutation, thus highlighting the homoplasy nature of Q675H spike mutation and the pivotal role of histidine residues near the furin cleavage site for SARS-CoV-2 adaptive evolution. We suggest that the RBD of the spike protein is a major hot spot region for SARS-CoV-2 evolution, while the furin cleavage site represents a secondary spot of adaptation to the human host. Based on this assumption, our data suggest that whenever a new variant appears carrying new mutations in its RBD domain, it replaces the less adapted one, which gradually disap-pears. Then, the new variant starts to acquire the Q675H mutation to gain a better fit-ness. This event occurs every time a better adapted variant appears.

This finding supports the hypothesis that VOC evolution aims not only at improving ACE2 affinity, but also at gaining more fitness to the human host, facilitating spike cleavage by furin. As a consequence, the number of VOC carrying Q675H spike is con-tinuously rising.

The functional characterization of Q675H in silico showed that this mutation, located in the P11 position of the polar region of the furin cleavage site, facilitates the accessi-bility of the core region to the furin binding pocket [28]. In particular, Q675H mutation forms an H-bonds network in the polar region delimiting the conformational space of the loop and improves its exposure to the protease. This implies an optimization of the directionality of the arginine residues involved in the interaction with the furin bind-ing pocket. Interestingly, this conformational restrain forces the peptide chain to change direction and separate the cleavage site from the other structural elements, thus resulting in a better exposure of cleavage site to the protease [50]. Our models show for the first time that P1-S1/P3-S4 substrate conformation puts the guanidine side chain of the R683 in position P3 in pocket S4, suggesting the P3 arginine as an im-portant residue for furin binding. It is worth noting that the P3 arginine is already known to be crucial for furin-dependent in vitro cleavage of a substrate construct car-rying the sequence of the S1/S2 site [11]. This new substrate conformation has been taken into consideration because the canonical furin recognition motif R-X-K/R-R*X (P4P3P2P1*P1’) does not contain an arginine residue in position P3, whereas in the SARS-CoV-2 spike the furin recognition motif is RRAR*S (P4P3P2P1*P1’) and displays arginine in position P3, which is close to P4 position in the three-dimensional space and can compete in the occupation of pocket S4. On the other hand, the Q675H mutant is the only one to have a P1-S1/P4-S4 conformation [51]. Importantly, the substrate conformations of Q675H have a higher binding affinity than the Q675 ones. This is due to the fact that Q675H always presents a y-conformation with the three arginine resi-dues alternatively arranged. This decreases the steric hindrance between them and consequently the arginine residues involved in the interaction can fit better in the binding pocket. On the whole, our analysis suggests that SARS-CoV-2 Q675H spike mutation enhances its availability to the furin proteolytic cleavage. As a result, Q675H mutation is likely to better promote spike S1 unit release and its interaction with hu-man receptor ACE2. In this study we predicted the effect of the Q675H mutation on the furin enzymatic activity. Other studies are necessary to evaluate how the Q675H influences the activity of the other proprotein convertases involved in spike activation.

The relevance of the furin cleavage site is highlighted by the significant number of mutations detected in this genomic region in different SARS-CoV-2 variants. Indeed, similarly to SARS-CoV-2 Q675H mutants, the emergence and rapid spread of a new variant, defined in Nexstrain as 20C-US, was documented in Louisiana and New Mex-ico (USA). This variant carries a series of mutations in SARS-CoV-2 genome, among which the spike substitution Q677H [52, 53], highlighting the crucial role of histidine in the release of SARS-CoV-2 S1 and S2 units operated by furin. Furthermore, B.1.617 lineage carries the P681R mutation in the furin cleavage site which increases S1/S2 cleavability, thus enhancing the viral fusion to the host cell membrane [54].

Notably, another mutation, the Q675K in the furin cleavage site has been reported, but its role in SARS-CoV-2 evolution still needs to be investigated [55, 56]. These findings further link spike mutation in the furin binding site to SARS-CoV-2 evolution in searching for the best fitness to increase its infectivity and spreading.

## 5. Conclusions

In conclusion, our data emphasize the role of the furin binding pocket for virus aggres-siveness and transmission, and highlight the importance of phylogenetic and biocom-putational analysis for better understanding SARS-CoV-2 evolutionary trajectory. A continuous genomic surveillance is crucial to deepen our understanding in SARS-CoV-2 evolution, especially in light of the ongoing vaccination campaign.

## Author Contributions

Conceptualization, P.D., M.C. and A.C.; methodology, A.B., G.C., S.M., M.M. and N.G.; formal analysis, A.B., G.C., S.M., M.M., M.G. and F.C.; investigation, A.B., G.C., S.M., M.M., M.G. and F.C.; writing – original draft preparation, A.B., P.D., M.C., F.C.; writ-ing—review and editing, A.C. and F.C. All authors have read and agreed to the published version of the manuscript.”

## Data Availability Statement

Genomic data reported in this study are available at Global Initiative on Sharing All Influenza Data (GISAID). Accession numbers: EPI_ISL_2671358, EPI_ISL_2671359, EPI_ISL_2671356, EPI_ISL_2671357, EPI_ISL_2671069, EPI_ISL_2671363, EPI_ISL_2671364, EPI_ISL_2671361, EPI_ISL_2671362, EPI_ISL_2671360, EPI_ISL_2665395.

## Acknowledgments

We acknowledge Spedali Civili of Brescia (Italy) for providing necessary in-frastructural support. We are grateful to the GISAID initiative and to its data contributors for making SARS-CoV-2 sequence data available in short time.

